# The Atypical Antipsychotic Quetiapine Induces Multiple Antibiotic Resistance in *Escherichia coli*

**DOI:** 10.1101/2021.11.29.470511

**Authors:** Yasuhiro Kyono, Lori Ellezian, YueYue Hu, Kanella Eliadis, Junlone Moy, Elizabeth B. Hirsch, Michael J. Federle, Stephanie A. Flowers

**Affiliations:** Department of Pharmacy Practice, College of Pharmacy, University of Illinois at Chicago; Department of Experimental and Clinical Pharmacology, College of Pharmacy, University of Minnesota; Department of Pharmaceutical Sciences, College of Pharmacy, University of Illinois at Chicago

## Abstract

Atypical antipsychotic (AAP) medication is a critical tool for treating symptoms of psychiatric disorders. While AAPs primarily target dopamine (D2) and serotonin (5HT2A and 5HT1A) receptors, they also exhibit intrinsic antimicrobial activity as an off-target effect. Because AAPs are often prescribed to patients for many years, a potential risk associated with long-term AAP use is the unintended emergence of bacteria with antimicrobial resistance (AMR). Here, we show that exposure to the AAP quetiapine at estimated gut concentrations promotes AMR in *Escherichia coli* after six weeks. Quetiapine-exposed isolates exhibited an increase in minimal inhibitory concentrations (MICs) for ampicillin, tetracycline, ceftriaxone, and levofloxacin. By whole genome sequencing analysis, we identified mutations in genes that confer AMR, including the repressor for the multiple antibiotic resistance *mar* operon (*marR*), and real-time RT-qPCR analysis showed increased levels of *marA, acrA*, and *tolC* mRNAs and a reduced level of *ompF* mRNA in the isolates carrying *marR* mutations. To determine the contribution of each *marR* mutation to AMR, we constructed isogenic strains carrying individual mutant *marR* alleles in the parent background and re-evaluated their resistant phenotypes using MIC and RT-qPCR assays. While *marR* mutations induced a robust activity of the *mar* operon, they resulted in only a modest increase in MICs. Interestingly, although these *marR* mutations did not fully recapitulate the AMR phenotype of the quetiapine-exposed isolates, we show that *marR* mutations promote growth fitness in the presence of quetiapine. Our findings revealed an important link between the use of AAPs and AMR development in *E. coli*.

## INTRODUCTION

The emergence of antibiotic-resistant bacteria is a worldwide threat to public health. In the US, the Centers for Disease Control and Prevention state that each year at least 2.8 million people experience infections caused by antibiotic-resistant bacteria, and more than 35,000 people die from organisms that have developed antimicrobial resistance (AMR) (1). While routine and widespread use of antibiotics is largely responsible for the acquisition of AMR among bacteria, increasing evidence suggests that non-antibiotics also play a role in AMR development due to their documented in vitro activity against microbes (2, 3).

An example of such drugs includes the class of antipsychotic compounds. Second-generation antipsychotics, often referred to as atypical antipsychotics (AAPs), are considered drugs of choice for treating individuals with schizophrenia (4). The use of AAPs has expanded to other Food and Drug Administration (FDA)-approved indications for other mental illnesses such as bipolar disorder, treatment-resistant depression, and autism-related irritability (5). It is estimated that approximately 1.7% of the US population is treated with an AAP drug (6). In recent years, AAPs have been widely prescribed to more vulnerable populations, such as in children, elderly, and pregnant women, in many cases for ‘off-label’ indications such as attention-deficit/hyperactivity disorders and disruptive behavior disorders (5, 7–9). This expansion of use has correlated with an increase in environmental detection of AAP drugs in municipal wastewater, hospital sewage, and drinking water (10–12).

Antimicrobial activity is well documented for older antipsychotics, and responses are reportedly similar to those evoked by sub-inhibitory concentrations of beta-lactam antibiotics (13–15). Studies in recent years have also shown that AAPs exhibit activity against specific commensal gut bacteria at estimated gut concentrations (2). AAPs are highly lipophilic and relatively poorly absorbed from the small intestine (16); therefore, substantial portions of these drugs travel through to the large intestine where most gut-dwelling bacteria reside. Atypical antipsychotics target dopamine and serotonin receptors in the brain that are generally absent in bacteria (17); thus, the mechanism by which AAPs exert antimicrobial activity to bacteria is still largely unknown. However, we and others have previously shown that AAP treatment is associated with measurable differences in the composition of gut microbial communities in preclinical models and psychiatric cohorts (18–20). Despite the known antimicrobial activity of AAP drugs, there is still a limited understanding on whether these drugs contribute to the emergence of AMR.

In this study, we investigated whether exposure to the AAP quetiapine results in the emergence of antibiotic resistance in the model bacteria, *Escherichia coli* –a commensal microbe in the human gut, which is also commonly present in drug-polluted environments. Quetiapine, a dibenzothiazepine class of AAPs, has seen a large increase in use since 2007 for both indicated and ‘off-label’ applications (21, 22). We exposed ATCC 25922 *E*.*coli* strain to two different doses of quetiapine *in vitro* for six weeks. For isolates exposed to quetiapine, we determined the minimal inhibitory concentration (MIC) to several different classes of antibiotics, and investigated the underlying AMR mechanism using whole-genome sequencing (WGS) and targeted gene expression analysis. These initial analyses led us to a discovery that quetiapine-exposed isolates acquired various mutations in the gene that encode a transcriptional repressor for the multiple antibiotic resistance (*mar*) operon whose activity is linked to AMR development. To further dissect the contribution of *marR* mutations to AMR development, we reconstructed *marR* mutant alleles in the parent *E*.*coli* strain background, and re-examined MIC, target gene expression analysis, and growth fitness. This work represents the first insight into a potential role of AAPs in contributing to AMR development by non-antibiotics.

## RESULTS

### Exposure to quetiapine results in antimicrobial resistance

The reference strain of *E. coli* (ATCC 25922) was serially passed every 24 hours in Luria Broth (LB) medium over 43 days. In parallel, *E. coli* was also passed in LB media containing either a ‘low-dose’(LD) quetiapine (10 µg/ml) and a ‘high-dose’(HD) quetiapine (100 µg/ml) over the same time period. All drug environments were performed in triplicate as biological replicates. To broadly screen for the development of AMR, mutation frequencies (MFs) were periodically calculated by plating all drug conditions on LB agar plates containing supra-MIC concentrations of selected antibiotics with differing mechanisms of action; (tetracycline [2 µg/ml], amikacin [16 µg/ml], and ampicillin [8 µg/ml]). By day 43, a modest but detectable change in MFs were calculated for both ampicillin (6.30E-07:LD and 2.10E-07:HD) and tetracycline (1.40E-09:LD and 4.10E-09:HD) antibiotics. Mutation frequencies for amikacin were unremarkable.

MICs were performed on a selection of isolates that grew on the antibiotic-containing agar plates to a panel of antibiotics including: ampicillin, amikacin, ceftriaxone, levofloxacin, and tetracycline. Tested isolates achieved at least a 16-fold increase in MIC dilutions to ampicillin (64 µg/ml), levofloxacin (0.5 µg/ml) and tetracycline (16 µg/ml) from the control background strain (**Table 1**). Isolates achieving the level of ‘clinical resistance’ were detected for ampicillin (64 µg/ml; HD only) and tetracycline (16 µg/ml; both LD and HD). We did not observe major differences between AMR development based on quetiapine doses. To evaluate the stability of antimicrobial resistance, isolates were then grown in quetiapine-free LB broth for 3 additional days and MICs were re-evaluated (**Table 1**). While resistance to ampicillin was stable after the discontinuation of quetiapine exposure, resistance to tetracycline was decreased by 2 to 4-fold, suggestive of transcriptional changes.

**Table 1:** Minimal inhibitory concentrations (MICs) of quetiapine-exposed isolates and reconstructed *marR* mutant strains

|  |  | Ampicillin (mg/L) |  |  | Amikacin (mg/L) |  |  | Ceftriaxone (mg/L) |  |  | Levofloxacin (mg/L) |  |  | Tetracycline (mg/L) |  |  |
| --- | --- | --- | --- | --- | --- | --- | --- | --- | --- | --- | --- | --- | --- | --- | --- | --- |
| marR genotype | Que | isolates | isolates (LB) | mutants | isolates | isolates (LB) | mutants | isolates | isolates (LB) | mutants | isolates | isolates (LB) | mutants | isolates | isolates (LB) | mutants |
| Parent (25922) |  | 4 |  |  | 16 |  |  | 0.06 |  |  | 0.03 |  |  | 1 |  |  |
| Isolate A (C108Y) | LD | 16 | 16 | 4 | 32 | 16 | 32 | 0.5 | 0.5 | 0.125 | 0.25 | 0.25 | 0.06 | 16 | 4 | 1 |
| Isolate B (A70fs) | LD | 32 | 32 | 8 | 16 | 32 | 32 | 0.5 | 0.5 | 0.25 | 0.25 | 0.25 | 0.06 | 16 | 4 | 2 |
| Isolate C (N126fs) | LD | 16 | 16 | 8 | 32 | 32 | 32 | 1 | 1 | 0.25 | 0.25 | 0.25 | 0.125 | 16 | 8 | 2 |
| Isolate D (L46P) | HD | 32 | 128 | 8 | 16 | 32 | 32 | 0.5 | 0.5 | 0.25 | 0.25 | 0.25 | 0.06 | 16 | 8 | 2 |
| Isolate E (A70fs) | HD | 64 | 128 | 8 | 32 | 32 | 32 | 1 | 1 | 0.25 | 0.5 | 0.25 | 0.06 | 16 | 4 | 2 |
| WT (isolate F); A70A ('mutant') | HD | 64 | 256 | 4 | 64 | 32 | 32 | 1 | 1 | 0.06 | 0.5 | 0.25 | 0.03 | 16 | 4 | 1 |

Que: Quetiapine; LD: Low Dose Quetiapine (10mg/L); HD: High Dose Quetiapine (100 mg/L); isolates: Isolates derived from quetiapine-exposed cultures after 43 days of exposure; isolates (LB): isolates that were grown in luria broth for three days without exposure to quetiapine; mutants: reconstructed strains that carry mutant *marR* alleles in the parent strain background

### Whole Genome Sequencing (WGS) Analysis Revealed Mutations in Antibiotic Resistant Genes

To identify mutations that may explain AMR phenotypes among the quetiapine-exposed isolates, we conducted WGS for genomic DNA isolated from: 1) Day three and Day 43 reference cultures grown in quetiapine-free environment; 2) two isolates (A,C) exposed to a low-dose quetiapine; and 3) two isolates (E,F) exposed to a high-dose quetiapine. The initial reference culture (Day 3) contained 67 genetic differences from the reference genome assembly of the ATCC 25922 strain, and the final culture (Day 43) additionally acquired 56 and lost six mutations between the two time points; we filtered out these mutations for further analysis. Among the four sequenced isolates, we identified a total of 291 unique mutations (**Supplemental Table 1**), nine of which were shared among all four, 27 were shared among three of four isolates (interestingly, all 27 were shared among isolates: A, C and E), and three were shared between different combinations of two isolates. Each isolate acquired 69 (A), 58 (C), 51 (E), and 74 (F) mutations that were not shared with others (**Figure 1**;(23)). In addition to these mutations, all four isolates shared a ∼52 kb deletion flanked by genes that are predicted to encode transposable elements (**Supplemental Table 2**). The deleted region contained 51 genes, including three annotated clusters of genes: 1) the *nanR* regulon that is involved in sialic acid transport and metabolism; 2) the *iucABCD-iutA* operon that is involved in aerobactin synthesis and transport; and 3) the *pap* operon that encodes the P fimbriae on the cell surface.

**Figure 1:**
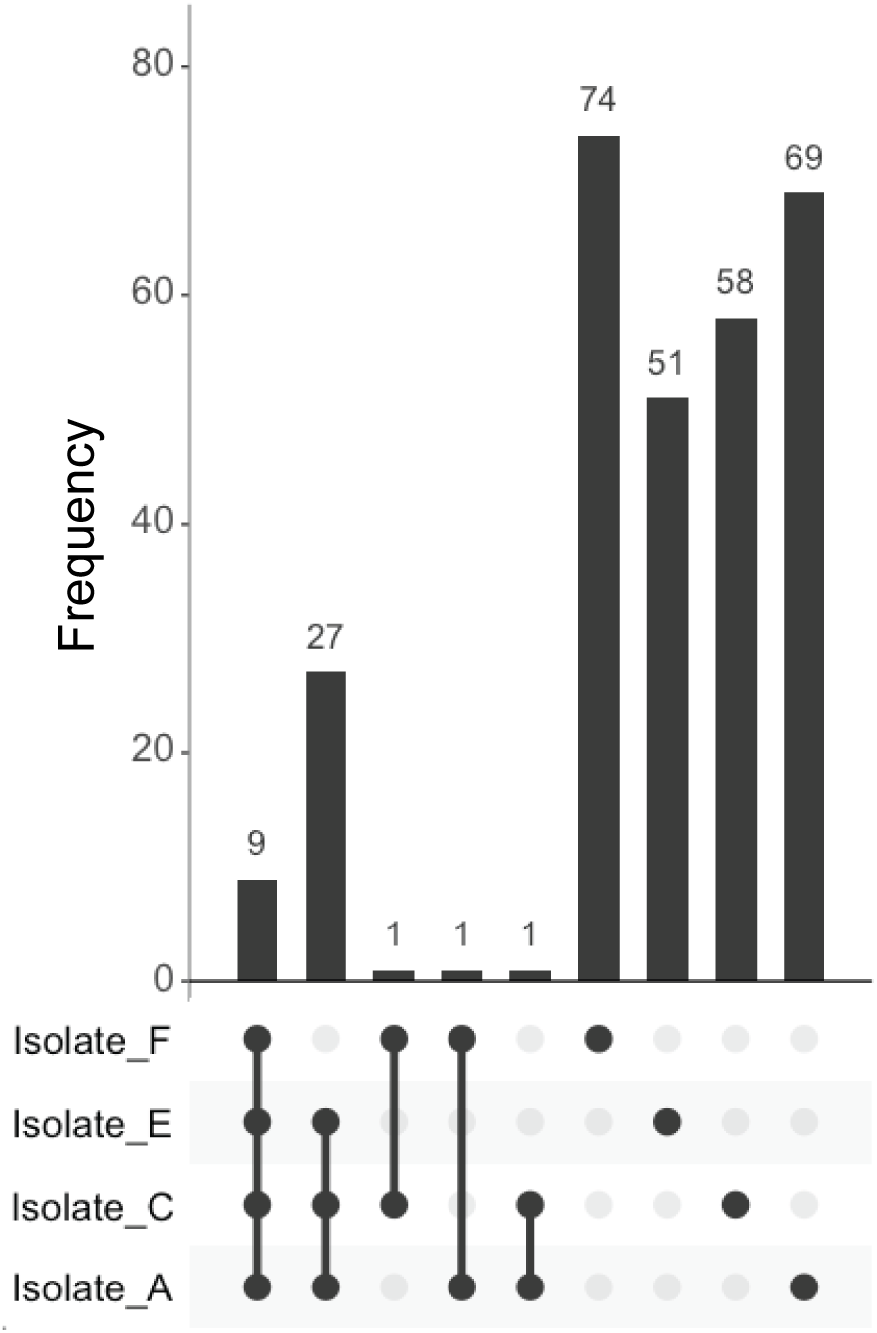
A summary of mutations identified by whole-genome sequencing analysis. The y-axis shows the number of mutations that were unique to each isolate or shared among multiple isolates. Combinations with a zero frequency were omitted from the plot.

We queried the comprehensive antibiotic resistant database (CARD) (24) to identify mutations in genes associated with AMR and found 14 gene matches (**Table 2**). From our sampled isolates, we observed 11 genes that carried either mutations that alter amino acid sequences (missense/nonsense mutations) or intergenic mutations that may alter its gene transcription (*acrB, acrE, acrF, acrR, arnT, basS (pmrB), ddlA, emrA, gspD, marR, msbA*); whereas, the rest carried a synonymous mutation (*baeS, glpT, gltB*). Two mutations were shared among the isolates: R497H missense mutation of the *msbA* gene (isolates A, C, and E), and various mutations of the *marR* gene (isolate A - C108Y missense mutation; isolate C - N126 frameshift mutation; and isolate E - A70 frameshift mutation). Other mutations were unique to each isolate and these are listed in **Table 2**.

**Table 2:**
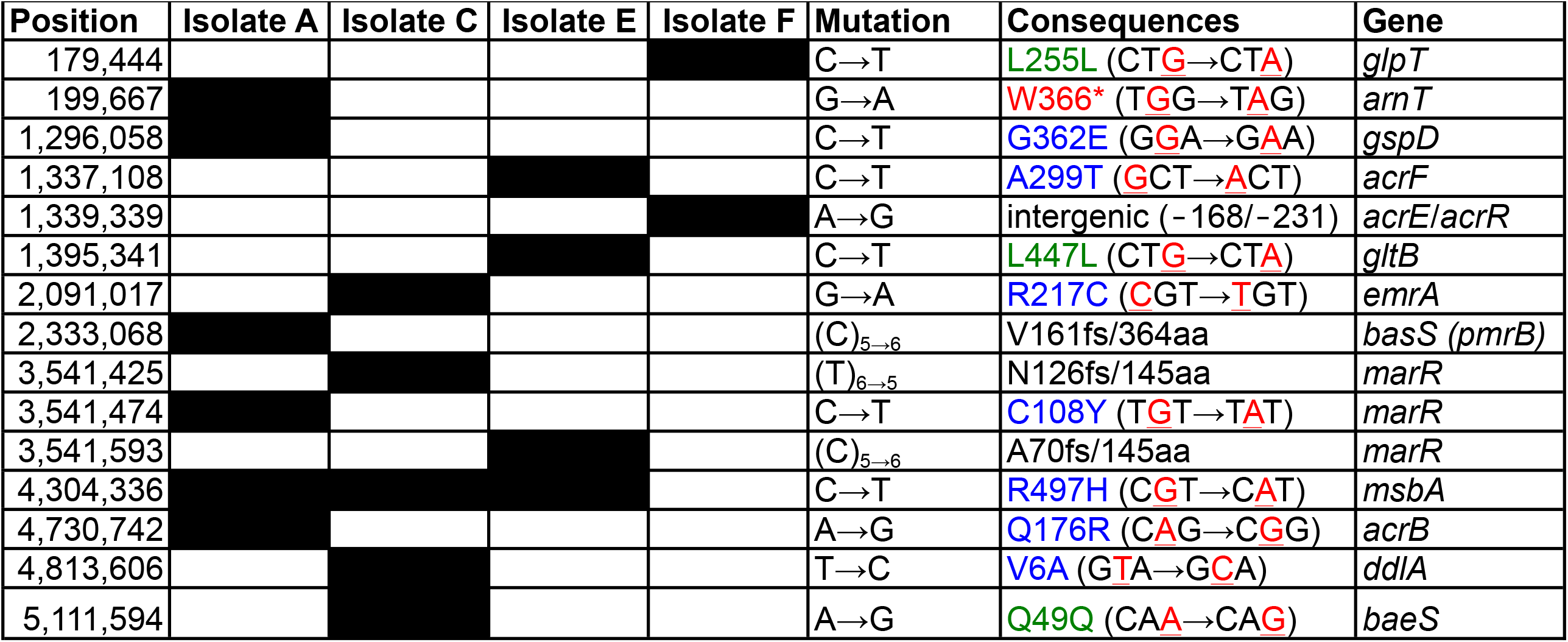
A list of chromosomal mutations identified in each isolate using whole-genome sequencing for genes that had entry matches in the CARD (filled boxes indicate the presence of mutations in isolates). Genome coordinate is based on the reference assembly of the ATCC 25922 strain (CP009072.1). *Intergenic* under the ‘Consequences’ column indicates mutations that were found in non-coding regions, and the numbers within the parentheses indicate the relative distance in base pairs from the closest flanking genes (Positive numbers indicate the distance from the translation termination site, while negative numbers indicate the distance from the translation start site). Frame shift (‘*fs’*) mutations are annotated by the first amino acid affected and its position, followed by the length of full protein.

The MarR protein is a direct transcriptional repressor for the *marRAB* operon. Loss-of-function mutations in the MarR protein leads to upregulation of *marA*, which in turn activates expression of genes that are involved in development of AMR, such as *acrA* and *tolC* genes that encode the AcrAB-TolC efflux pump (25). Indeed, three of the four sequenced isolates acquired mutations in the *marR* gene. We genotyped two additional isolates and found that both also contained mutations in the *marR* gene (isolate B - A70 frameshift mutation; and isolate D - L46P missense mutation). In total, five of the six quetiapine-exposed isolates have acquired a mutation in the *marR* gene (isolates A-E) (**Figure 2**); and therefore, for the remainder of the study we investigated the functional relevance of these *marR* mutations to AMR induced by quetiapine exposure.

**Figure 2:**
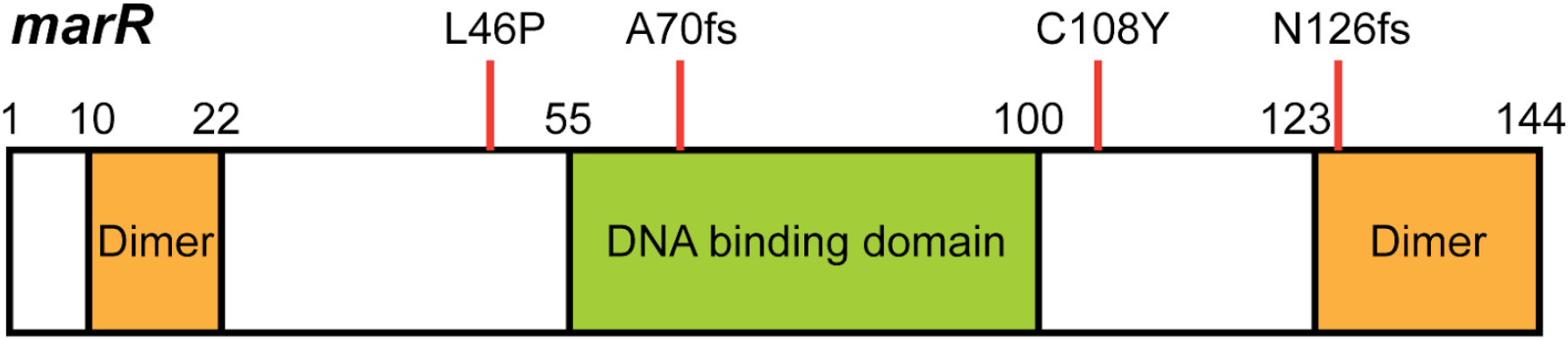
Location of mutations found within the *marR* gene among quetiapine-exposed isolates.

### Quetiapine-exposed isolates exhibit increased levels of *marA, acrA, and tolC* mRNA and reduced level of *ompF* mRNA

We conducted real-time RT-qPCR analysis for representative MarR-regulated genes: 1) *marA* transcriptional activator within the *marRAB* operon that is directly repressed by MarR (primary target); 2) *acrA* and *tolC* genes that are directly activated by MarA (secondary target); and 3) *ompF* porin gene that MarA indirectly downregulates via activation of *micF* (tertiary target) (26). Compared to the parent WT strain, quetiapine-exposed isolates with a *marR* mutation (isolates A-E) expressed higher mRNA levels of *marA* (92.1-fold increase on average), *acrA* (4.0-fold increase), and *tolC* genes (4.5-fold increase) and a lower level of *ompF* mRNA (14.3-fold decrease) (**Table 3**). Isolate F did not carry a *marR* mutation; however, it also exhibited a modest increase in mRNA levels of *marA* (2.4-fold increase) and *acrA* genes (1.9-fold increase), while the level of *ompF* mRNA showed a similar degree of decrease compared to the other isolates (5.9-fold decrease); *tolC* mRNA did not exhibit a statistically significant change (1.9-fold increase, *p*=0.055). These results indicate that *marR* mutations identified in quetiapine-exposed isolates led to a robust activation of the *marRAB* operon.

**Table 3:** Summary of real-time RT-qPCR analysis in quetiapine-exposed isolates. Shown are means and SEM of the normalized mRNA levels of *marA, acrA, tolC* and *ompF* genes between two replicates. Asterisks indicate statistical significance compared to the parent WT strain (p < 0.05). Fold-change (FC) is relative to the normalized quantity of the parent WT strain.

| *marR* genotype | Que | *marA* |  |  |  | *acrA* |  |  |  |
| --- | --- | --- | --- | --- | --- | --- | --- | --- | --- |
|  |  | Isolates | FC | Isolates (LB) | FC | Isolates | FC | Isolates (LB) | FC |
| 25922 (WT) |  | 0.045 ± 0.003 | - | - | - | 0.494 ± 0.003 | - | - | - |
| Isolate A (C108Y) | LD | 1.971 ± 0.167* | 43.5 | 2.010 ± 0.133* | 44.4 | 2.822 ± 0.184* | 5.7 | 1.587 ± 0.127* | 3.2 |
| Isolate B (A70fs) | LD | 4.464 ± 0.313* | 98.5 | 3.450 ± 0.055* | 76.1 | 1.547 ± 0.111* | 3.1 | 1.674 ± 0.073* | 3.4 |
| Isolate C (N126fs) | LD | 6.196 ± 0.859* | 136.7 | 6.961 ± 0.178* | 153.6 | 2.240 ± 0.269* | 4.5 | 1.908 ± 0.201* | 3.9 |
| Isolate D (L46P) | HD | 4.662 ± 0.126* | 102.9 | 1.100 ± 0.088* | 24.3 | 1.509 ± 0.009* | 3.1 | 3.776 ± 0.254* | 7.6 |
| Isolate E (A70fs) | HD | 3.583 ± 0.063* | 79.1 | 2.525 ± 0.070* | 55.7 | 1.663 ± 0.008* | 3.4 | 2.003 ± 0.103* | 4.1 |
| Isolate F (WT) | HD | 0.107 ± 0.008* | 2.4 | 0.060 ± 0.003 | 1.3 | 0.948 ± 0.070* | 1.9 | 0.705 ± 0.048* | 1.4 |
| *marR* genotype | Que | *tolC* |  |  |  | *ompF* |  |  |  |
|  |  | Isolates | FC | Isolates (LB) | FC | Isolates | FC | Isolates (LB) | FC |
| 25922 (WT) |  | 0.577 ± 0.005 | - | - | - | 4.733 ± 0.180 | - | - | - |
| Isolate A (C108Y) | LD | 2.547 ± 0.044* | 4.4 | 2.168 ± 0.052* | 3.8 | 0.956 ± 0.004* | 0.20 | 0.586 ± 0.001* | 0.1 |
| Isolate B (A70fs) | LD | 2.064 ± 0.113* | 3.6 | 2.045 ± 0.029* | 3.5 | 0.057 ± 0.000* | 0.01 | 0.265 ± 0.009* | 0.1 |
| Isolate C (N126fs) | LD | 3.598 ± 0.431* | 6.2 | 2.320 ± 0.132* | 4 | 0.134 ± 0.005* | 0.03 | 0.327 ± 0.004* | 0.1 |
| Isolate D (L46P) | HD | 2.412 ± 0.056* | 4.2 | 1.463 ± 0.068* | 2.5 | 0.046 ± 0.003* | 0.01 | 0.719 ± 0.002* | 0.2 |
| Isolate E (A70fs) | HD | 2.448 ± 0.060* | 4.2 | 1.889 ± 0.003* | 3.3 | 0.545 ± 0.004* | 0.12 | 0.568 ± 0.016* | 0.1 |
| Isolate F (WT) | HD | 1.102 ± 0.128 | 1.9 | 0.652 ± 0.027 | 1.1 | 0.788 ± 0.025* | 0.17 | 4.477 ± 0.032 | 1.0 |

To examine the stability of these transcriptional changes, we cultured these isolates in quetiapine-free growth media for three additional days, and re-evaluated mRNA levels using RT-qPCR. Compared to the parent WT strain, isolates A-E maintained higher mRNA levels of *marA* (70.8-fold increase on average), *acrA* (4.4-fold increase), and *tolC* genes (3.4-fold increase), and a lower mRNA level for the *ompF* gene (10-fold decrease on average). Meanwhile, mRNA levels in isolate F had decreased to the levels that were comparable to those of the parent WT strain (*marA:* 1.3-fold; *acrA:* 1.4-fold, *tolC:* 1.1-fold, and *ompF:* 0.95-fold, respectively; **Table 3**).

### Variant *marR* alleles only modestly contribute to antimicrobial resistance despite a robust increase in *marRAB* operon activity

We hypothesized that *marR* mutations contributed to the reduced antimicrobial susceptibility among quetiapine-exposed isolates. To test the hypothesis, we reconstructed isogenic strains that carry mutant *marR* alleles (C108Y, A70fs, N126fs, and L46P) in the ATCC 25922 strain background using lamda red recombineering. As acontrol, we also constructed a mock mutant strain that carried a A70A synonymous mutation (GC<u>A</u>→ GC<u>G</u>).

We first conducted real-time RT-qPCR for the same subset of *marRAB* operon genes (*marA, acrA, tolC* and *ompF*) to determine the level of *marRAB* operon activity in these reconstructed strains. All four genes exhibited the same direction of changes in mRNA levels as quetiapine-exposed isolates with the matching *marR* genotypes (*marA*: 39.3-fold increase on average; *acrA*: 1.6-fold increase, *tolC*: 2.6-fold increase; and *ompF*: 6.7-fold decrease); however, the degree of change was not as robust as those of matching isolates (**Table 4**). The A70A control strain did not show significant changes in mRNA levels of any target genes.

**Table 4:** Summary of real-time RT-qPCR analysis in reconstructed strains with mutant *marR* alleles. Shown are means and SEM of the normalized mRNA levels of *marA, acrA, tolC* and *ompF* genes between two replicates. Asterisks indicate statistical significance compared to the parent WT strain (*p* < 0.05). Fold-change (FC) is relative to the normalized quantity of the parent WT strain.

| *marR* genotype | Corresponding isolates | *marA* |  | *acrA* |  | *tolC* |  | *ompF* |  |
| --- | --- | --- | --- | --- | --- | --- | --- | --- | --- |
|  |  | Mutants | FC | Mutants | FC | Mutants | FC | Mutants | FC |
| 25922 (WT) | - | 0.037 ± 0.006 | - | 0.608 ± 0.027 | - | 0.405 ± 0.009 | - | 2.466 ± 0.197 | - |
| SF003 (C108Y) | A | 0.603 ± 0.076* | 16.5 | 0.723 ± 0.096 | 1.2 | 0.786 ± 0.113 | 1.9 | 0.455 ± 0.034* | 0.2 |
| SF004 (A70fs) | B,E | 1.394 ± 0.144* | 38.0 | 0.991 ± 0.090* | 1.6 | 1.073 ± 0.036* | 2.7 | 0.357 ± 0.056* | 0.1 |
| SF005 (N126fs) | C | 2.294 ± 0.254* | 62.6 | 1.183 ± 0.071* | 1.9 | 1.235 ± 0.044* | 3.1 | 0.281 ± 0.018* | 0.1 |
| SF006 (L46P) | D | 1.513 ± 0.163* | 41.3 | 1.081 ± 0.081* | 1.8 | 1.087 ± 0.046* | 2.7 | 0.356 ± 0.026* | 0.1 |
| SF007 (A70A) | F | 0.035 ± 0.003 | 1.0 | 0.563 ± 0.043 | 0.9 | 0.425 ± 0.041 | 1.0 | 2.459 ± 0.290 | 1.0 |

We conducted MICs against the same panel of antibiotics (ampicillin, amikacin, ceftriaxone, levofloxacin, and tetracycline) and evaluated changes in MICs relative to the parent WT strain. For ampicillin, amikacin, and tetracycline, the reconstructed strains showed either no change, or only two-fold increase in MICs (**Table 1**). For ceftriaxone, A70fs, N126fs and L46P alleles exhibited four-fold increase in MIC, while the C108Y allele only exhibited a two-fold increase. For levofloxacin, the N126fs allele resulted in a four-fold increase in MIC, while the other three showed a two-fold increase. We did not observe any significant changes in MICs for all five antibiotics for the control A70A mutant.

### Relative fitness of *marR* mutations increases as a function of quetiapine concentration

Despite the robust activation of the *marRAB* operon, the mutant *marR* alleles only conferred a modest, if any, increase in antimicrobial MICs. As published work has shown that mutations in the *marR* gene confer growth advantage in the presence of ciprofloxacin (27), we hypothesized that mutant m*arR* alleles might confer growth advantage in the presence of quetiapine. We tested the hypothesis by conducting a growth curve analysis in media containing increasing concentrations of quetiapine (0, 100, 300, and 500 g/mL) (**Figure 3; Supplemental Figure 1**). We measured two fitness components: 1) growth rate (r), and 2) the maximum population size at the stationary phase, also termed the carrying capacity (K). We found a significant dose-dependent effect of quetiapine on the growth rate of parent WT strain (*p* = 1.97E-05; one-way ANOVA, followed by Fisher’s LSD *post hoc* test) but not on those for the mutant strains (**Figure 3, top panel)**. When growth rates were compared at each dose, the *marR* mutants grew significantly slower than the parent WT strain in quetiapine-free media (*p* = 0.023). Growth rates were similar between the parent and *marR* mutants in media containing 100 and 300 µg/mL quetiapine (*p =* 0.957 and *p =* 0.418, respectively). *MarR* mutants grew significantly faster at 500 µg/mL (*p* = 0.002), likely due to a significant decline in the growth rate of the parent WT strain, as this dose approaches MIC for quetiapine at 512 µg/mL (data not shown). The A70A control strain behaved similarly to the parent WT strain, except that it grew slower than the parent WT strain in the quetiapine-free media (*p* = 0.003; **Supplemental Figure 1**). The carrying capacity did not significantly differ among the WT parent or *marR* mutants at 0, 100, and 300 µg/mL doses of quetiapine (**Figure 3 - panel B)**; whereas at a 500 µg/mL dose, it was significantly diminished for both the WT parent and *marR* mutants (WT: *p* = 4.08E-04; mutants: *p* = 2.55E-06; one-way ANOVA, followed by Fisher’s LSD *post hoc* test). However, at the 500 µg/mL dose, *marR* mutants exhibited a significantly greater carrying capacity with respect to the WT parent strain (*p* = 0.001). These findings together support our hypothesis that *marR* mutations derived from quetiapine exposure confer a selective growth advantage in the presence of quetiapine. The growth fitness of the quetiapine-exposed isolates showed variability in both growth rate and carrying capacity, which is likely because these isolates carried other genetic alterations that might affect their growth pattern; however, we observed a consistent pattern of growth advantage in the presence of quetiapine for all six isolates, especially at the near-MIC dose compared to the parent WT strain (**Supplemental Figure 1**).

**Figure 3:**
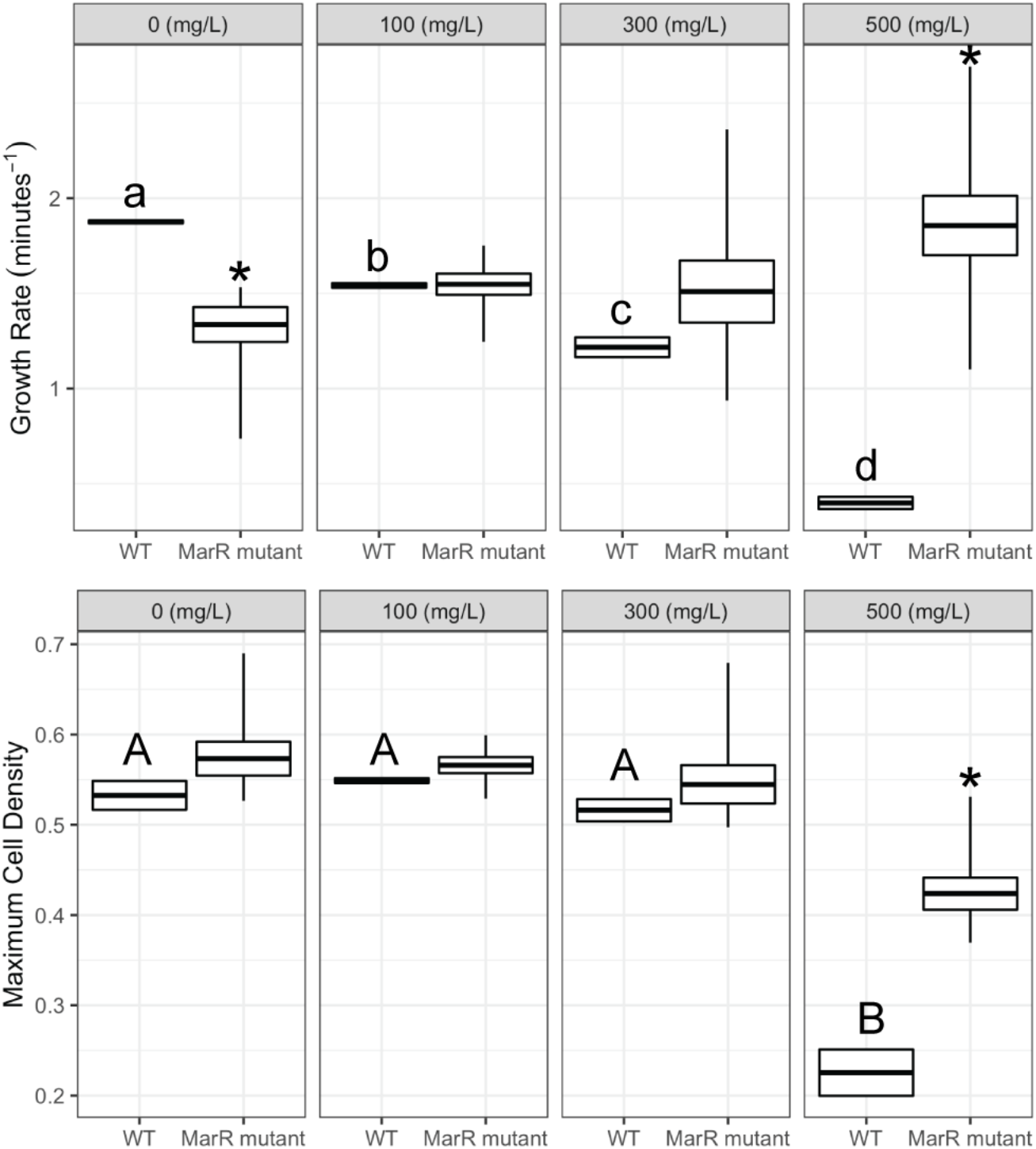
Growth curve analysis of the parent WT strain (*marR*^*WT*^) and reconstructed *marR* mutant strains grown in media containing 0, 100, 300, and 500 µg/mL quetiapine for 16 hrs. The top panels show growth rate (r), and the bottom panels show the maximum population at the stationary phase (carrying capacity, K). Box plots represent the mean ± SEM (duplicates), and WT means with the same letter (a, b, c for growth rate; A, B for the carrying capacity) are not significantly different (growth rate: *p* = 1.97E-05; carrying capacity: *p* = 4.08E-04; one-way ANOVA, followed by Fisher’s LSD *post hoc* test). Asterisks above the means of mutant strains indicate statistically significant differences in r/K compared to the WT strain treated with the same dose of quetiapine (growth rate at 0 µg/mL: *p =* 0.023; growth rate at 500 µg/mL: *p =* 0.002; carrying capacity at 500 µg/mL: *p* = 0.002; two-tailed Student’s *t* test).

## DISCUSSION

There is growing evidence that non-antibiotics contribute to the development of AMR in bacteria (2, 3). In this study, we discovered that exposure to the AAP quetiapine at estimated gut concentrations could give rise to multidrug AMR in *E. coli*. WGS analysis revealed that these antibiotic-resistant isolates carried mutations in genes that are known to induce AMR, including the *marR* gene, which we observed in five of six sequenced isolates. To investigate a contribution of *marR* mutations to AMR phenotype, we reconstructed isogenic strains with mutant alleles for each *marR* mutation observed among the isolates. Contrary to our hypothesis, these strains did not exhibit a robust AMR phenotype like the quetiapine-exposed isolates; however, *marR* mutations conferred a growth advantage for the strains in the presence of quetiapine in their growing environment. To our knowledge, this is the first known report that exposure to an AAP drug induces AMR in bacteria.

Mechanistic studies have identified the *marRAB* operon as an important contributor to fluoroquinolone resistance in uropathogenic *E. coli* (UPEC) and a virulence regulator in *E. coli* in a murine model of pyelonephritis (28). Previous work investigating *E. coli* with *marR* mutations have demonstrated a positive correlation between the level of *marA* expression and ciprofloxacin resistance (27). We made a similar observation that the *marR* mutation that induced the highest level of *marA* mRNA among the four mutations investigated in this study (N126fs) exhibited the most resistant phenotype with a four-fold increase in both ceftriaxone and levofloxacin MICs. However, the degree of resistance achieved by the reconstructed mutant *marR* strains was consistently less than that observed for the quetiapine-exposed isolates with matching *marR* genotypes. This is potentially due to a lower activity of *marRAB* operon genes, approximately 3-fold lower on average, compared to the level of activity within corresponding quetiapine-exposed isolates.

Individual mutations in resistance genes typically confer only an incremental increase in antimicrobial resistance in *E. coli* (29, 30). Therefore, in order to reach clinically relevant levels of resistance, *E. coli* may need to acquire several step-wise genetic alterations. Besides *marR*, our WGS analysis of the quetiapine-exposed isolates uncovered mutations in other genes that are implicated for AMR. Three of the isolates (A, E, and F) had mutations in members of the *acr* genes that belong to the resistance-nodulation-cell division (RND) antibiotic efflux pump, which is known to play a crucial role in intrinsic resistance to antibiotics in *E*.*coli*. First, isolate A had a missense mutation (Q176R) in the *acrB* gene, the inner membrane component of the AcrAB-TolC efflux pump. The Q176 residue aligns the wall of the drug binding pocket for the AcrB protein that regulates substrate specificity (31); therefore, this mutation may potentially alter AcrB function, as has been shown that another single amino acid substitution of this protein (G288D) led to altered drug specificity and increased efflux activity (32). Second, isolate E carried a missense mutation (A299T) in the *acrF* gene, which is the inner membrane component for another RND antibiotic efflux pump, AcrEF-TolC, that is predicted to function similarly to acrB (ATCC 25922’s AcrB and AcrF proteins share 77% amino acid sequence identity;(33)). While A299 residue is conserved between AcrB and AcrF proteins, it does not fall within any known functional domains in either of these two proteins, so functional relevance of this particular mutation is unclear. However, a prior study has shown that deletion of the *acrF* gene could lead to increased expression of other efflux pump genes such as *acrB* as a potential form of compensation (34); therefore, if A299T is a loss-of-function mutation, it may indirectly promote AMR phenotypes through such a mechanism. Lastly, isolate F also carried a mutation in the *acrEF* operon; unlike isolate E, however, isolate F has an intergenic mutation between the *acrE* and *acrR* genes, which happens to overlap with the -10 consensus sequence within the *acrE* promoter (TAGAA<u>T</u> → TAGAA<u>C</u>). Prior studies have shown that intergenic mutations that alter gene transcription can be the underlying mechanisms by which organisms acquire AMR phenotypes (35). Another AMR gene that was shared among quetiapine-exposed isolates (A, C, and E) was a missense mutation R497H in the *msbA* gene, which encodes ATP-binding cassette (ABC) transporters. R497 residue is located within a highly conserved ABC transporter domain; hence, the mutation has a potential to alter characteristics of its protein. While this particular mutation was not directly analyzed, mutations within this domain influences cell growth and the activity level of ATP hydrolysis (36).

While we did not observe a significant role for *marR* mutations in antimicrobial resistance, we demonstrated that these mutations were beneficial for growth in the environment in which quetiapine is present. As has been shown in previous work, *marR* mutations are energetically costly, and the presence of a *marR* mutation can decrease fitness without a selective pressure (37). Indeed, *marR* variants detected clinically in UPEC isolates often include compensatory mutations that mitigate this growth fitness cost but consequently attenuates antimicrobial resistance (37). Likewise, we observed a significant decrease in growth rate of *marR* mutant strains when compared to the parent WT strain in a quetiapine-free environment. However, this growth deficit was eliminated in the presence of quetiapine and became beneficial as the concentration of quetiapine approached the MIC for the parent. *MarR* has been shown to detect liberated copper from the cell membrane, explaining activation of the *marRAB* pathway with bactericidal antibiotics (38). Interestingly, published data suggest that antipsychotic drugs, including quetiapine, target cellular membranes leading to increased permeability in mammalian and Cryptococcus cell models (39, 40). Further studies are needed to understand the mechanisms by which quetiapine exposure leads to AMR development. Also, it warrants further investigation to determine whether the emergence of AMR is unique to quetiapine exposure, or if it is broadly applicable to other AAP agents.

Unlike antimicrobials, AAP medications are often indicated for long-term treatment periods, and therefore might represent a significant and unrecognized risk factor for the development of AMR in this patient population. Serious psychiatric disorders like schizophrenia are associated with increased risk of infections (41); however, the relationship among a psychiatric comorbidity, AAP treatment, and the frequency or severity of an infection is complex. The use of AAPs has been associated with a higher incidence of urinary tract infections (UTIs) in elderly and female patients and among patients with Parkinson’s disease treated with AAPs (42, 43). Furthermore, AAP treatment is an established risk factor for pneumonia and increased mortality from pneumonia in both hospitalized and elderly patients (44, 45). It is critical that we investigate how changes in gut bacteria in response to AAP exposure contributes to the efficacy and potential adverse effects of these medications, including the development of AMR in treated populations.

## MATERIALS AND METHODS

### Test Compound

The second-generation antipsychotic drug quetiapine fumarate (catalog #PHR1856) was manufactured by Sigma-Aldrich (St. Louis, MO).

### Antibiotics

The following antibiotics were purchased from Sigma-Aldrich: amikacin disulfate (catalog #A1774), ceftriaxone disodium salt hemi(heptahydrate) (#C5793), levofloxacin (#28266), and tetracycline hydrochloride (#T7660). Ampicillin sodium (cat #A-301-5) was purchased from Gold Biotechnology (St. Louis, MO).

### Bacterial subculturing and mutation frequency determination

*E. coli* (ATCC 25922) was serially passed every 24 hours in culture medium (LB: 10 g/L tryptone, 5 g/L yeast extract, 10 g/L NaCl) over 43 days. In parallel, *E. coli* was also passed in LB media containing either a ‘low-dose’(LD) quetiapine (10 µg/ml) and a ‘high-dose’(HD) quetiapine (100 µg/ml) over the same time period. Colonic quetiapine concentrations were estimated based on 1) average daily dose of quetiapine (400-800 mg/day); 2) percentage of dose reaching colon (46); 3) measurements of average colon fluid volumes (47); and 4) measurements of average adult total colon volume. All drug environments were performed in triplicate as biological replicates. Mutation frequencies (MFs) were calculated by plating all drug conditions on LB agar plates containing tetracycline (2 µg/mL), amikacin (32 µg/mL), or ampicillin (8 µg/mL). The mutation frequency was defined as the number of colonies grown on the antibiotic-infused agar plates divided by the total bacterial count.

### Minimal inhibitory concentrations

Minimal inhibitory concentrations (MICs) of antibiotic-resistant colonies were determined by broth microdilution method to ampicillin, amikacin, ceftriaxone, levofloxacin, and tetracycline, as recommended by CLSI (49). A bacterial sample was taken from cultured Petri dishes and diluted in 0.9% normal saline. The turbidity of each tube was adjusted to McFarland 0.5 scale (∼1.5 × 10^8^ CFU/mL), diluted 1/100 in appropriate volume of media, and then 100 µL of broth was added to wells 2-12 of a 96-well plate. Antibiotic stock, 200 µL, was then added to well 1. One hundred µL of drug stock was taken from well 1, and serially diluted from wells 2-12. One hundred µL of the bacterial dilution was added to each well except the negative controls, which only contained broth. This procedure was done in triplicate for each isolate or strain. The MIC was defined as the lowest concentration at which no microbial growth was observed after 24 hours of incubation.

### DNA extraction, whole-genome sequencing, and data analysis

Genomic DNA from overnight cultures was extracted using the DNeasy® Blood & Tissue Extraction Kit (Qiagen, Germantown, MD; #69504) according to the manufacturer’s instructions. The samples were prepared for sequencing by an initial quantification using Qubit 4 Fluorometer (Life Technologies, Grand Island, NY; #Q32851). Library preparation was performed using the Illumina DNA Prep Workflow with UDI indexing (Illumina Inc., San Diego, CA; #20018705, 20027213) according to the manufacturer’s instructions with 100 ng template input and 5 cycles of PCR. An equal-volume pool of all libraries was then made. The pool was quantified using a Qubit DNA High Sensitivity kit (Life Technologies, #Q32851, Grand Island, NY), and size distribution was assessed using an Agilent 4200 TapeStation System (Agilent Technologies, Santa Clara, CA; #G2991AA) using TapeStation D5000 ScreenTape, ladder and assay (Agilent Technologies, # 5067-5588, 5067-5590 and 5067-5589). The pooled libraries were run on the Illumina MiniSeq instrument using MiniSeq Reagent MO Kit (300 cycles) (Illumina Inc.) for quality control and library balancing purposes. A new pool was made based on the MiniSeq run results, quantified the same as described above and sequenced on an Illumina NovaSeq 6000 instrument (300 cycles) (Illumina Inc.), with a 1% phiX spike-in. We used the breseq computational pipeline to identify mutations (version 0.35.5;(50)) using the reference assembly of the ATCC 25922 strain (NCBI: CP009072.1) to align the Illumina reads. Because the coverage for each sample exceeded 290X, we downsampled coverage to 60X for AMR isolates (*-l 60*) and to 150X for reference culture samples (*-l 150*).

### Reconstruction of *marR* mutant strains

We reconstructed two independent strains for each *marR* mutation in the chromosomal locus of the ATCC 25922 *E*.*coli* reference strain using the selection-counter selection method facilitated by Lambda Red recombineering ((51), **Supplemental Table 3**). We first electroporated the pKD46 Lambda Red expression plasmid (52) into the reference strain using Bio-Rad Gene Pulser (2.5 kV, 200 ohm, 25 µF), and then selected for colonies on LB agar plates that contained 100 mg/mL ampicillin (LB-Amp). From this point forward, we maintained cultures/plates at 30°C to preserve the pKD46 plasmid.

#### First recombination

Cultures were grown overnight in LB-Amp liquid media. Next day, we transferred 1/100 volume of overnight culture to fresh LB-Amp media that additionally contained 1% L-arabinose (w/v) (Sigma-Aldrich #A3256) to induce the expression of Lambda Red recombinase, and shake-incubated it at 30°C until the culture reached McFarland 2 scale (equivalent of OD_600_ ∼0.6-0.8). Cells were made electro-competent by previously published methods (53). Briefly, we washed cultures twice with sterile water at room temperature, and after the final wash, resuspended cells in sterile water with 1/100 volume of the initial culture (e.g. 30 µL final cell suspension from 3 mL initial culture). We electroporated 25 µL of cell suspension with 100 ng of PCR-generated DNA fragments that encoded a synthetic *cat-sacB* expression cassette (GenBank: KM018298.1), and recovered the cells in SOC medium at 30°C for 2 hrs with vigorous shaking. When the cassette is successfully incorporated, the strain becomes both chloramphenicol-resistant and sucrose-sensitive. After recovery, cells were plated on LB-Lennox agar plates (10 g/L tryptone, 5 g/L yeast extract, 5 g/L NaCl) that contained 10 µg/mL chloramphenicol to select for successful recombinants, which was verified by Sanger sequencing.

#### Second recombination

We exchanged the *cat-sacB* cassette with DNA fragments that encoded mutant *marR*, which were PCR-generated from quetiapine-exposed isolates using their genomic DNA as PCR templates. Electroporation reactions were conducted as described above using 100 ng of PCR-amplified DNA fragments as substrates. Cells were recovered in no-salt growth media (10 g/L tryptone, 5 g/L yeast extract) for at least 4 hours to ensure the loss of SacB protein. After recovery, we counter-selected recombinants on no-salt LB agar plates with 5% sucrose (w/v) (10 g/L tryptone, 5 g/L yeast extract, 50 g/L D-sucrose). Sanger sequencing verified seamless incorporation of mutations at the chromosomal *marR* locus. To evaluate if our experimental procedures independently generated strains with antimicrobial resistance, we constructed two independent control strains (SF007a, SF007b) that carried a synonymous mutation (GC<u>A</u>→ GC<u>G</u>; A70A) in the *marR* gene. These control strains underwent the same experimental procedures as other reconstructed strains, except during the second recombination step, cells were electroporated with 1 µg of 89-bp oligonucleotides as a substrate to replace the *cat-sacB* cassette (oligonucleotide sequence is listed in **Supplemental Table 4)**

### Gene expression analysis by real-time RT-qPCR

A 1/100 dilution of overnight cultures was transferred to fresh media and shake-incubated until the turbidity of each tube reached McFarland 2 scale (∼6 × 10^8^ cfu/mL). We harvested 1 mL of each sample, incubated in RNAprotect® Bacteria Reagent (Qiagen, Germantown, MD) according to the manufacturer’s instructions, and stored at - 80°C until total RNA extraction using TRIzol™ Reagent (Invitrogen, Waltham, MA; #15596-018). We first treated 1 µg of extracted RNA with DNaseI (Invitrogen, #18068-015) to remove genomic DNA contamination, and then synthesized first-strand cDNA in the presence of RNAse inhibitor using the High Capacity cDNA Reverse Transcription Kit (Applied Biosystems, Waltham, MA; #4368814), during which we generated No-RT control samples to which no reverse transcriptase was added. We conducted quantitative real-time PCR on ViiA7 Real-Time PCR System with QuantStudio software (v1.3) (Applied Biosystems) using PowerUP SYBR Green Master Mix (Applied Biosystems, #A25742) with a fast cycling mode (1 cycle of 50°C for 2 min; 1 cycle of 95°C for 2 min; 40 cycles of 95°C for 1 second and 60°C for 30 seconds). To measure mRNA levels for each target gene (*marA, acrA, tolC*, and *ompF*), we constructed standard curves and normalized it to the mRNA level of *hcaT* housekeeping gene that did not exhibit significant changes among samples. The No-RT control samples did not generate a significant quantity of amplicons for any targets. Data are presented as means ± SEM, performed in duplicate. Primer sequences are listed in **Supplemental Table 4**.

### Growth curve analysis

We grew fresh quetiapine-exposed isolates on LB plates that contained 10 µg/mL quetiapine, and the reference ATCC 25922 strain or reconstructed *marR* mutant strains on LB plates without quetiapine. The next day, we resuspended colonies in 0.9% normal saline, adjusted the concentration of each sample to ∼3.0 × 10^8^ cfu/mL, and cultured 1/100 of each isolate/strain (in duplicates) in a sealed 96-well flat-bottom plate with LB media containing 0, 100, 300, or 500 µg/mL quetiapine (200 µL final volume in each well). The plate was shaken continuously at 37°C in the Synergy HTX multi-mode reader (BioTek Instruments; Winooski, VT), and OD_600_ was recorded every hour for 16 hrs. Reported OD_600_ were used to construct a growth curve, and the comparison of growth curve fitness components were calculated using the Growthcurver package for R (54).

### Statistical Analysis

Statistics were performed using R v3.2.5 (R Foundation for Statistical Computing, Vienna, Austria). Differences in resistance or fitness phenotype between treatment groups were determined by two-tailed Student’s *t* tests (ɑ = 0.05). One-way ANOVA followed by Fisher’s least significant differences (LSD) *post hoc* test was used for multiple comparisons.

## Data Availability

Raw sequencing is available on the NCBI Sequence Read Archive under accession no. XXXXXXXXX.

## Acknowledgments

L.E. was supported by the Riback Fellowship at the University of Illinois at Chicago. We would like to thank Dr. Diarmaid Hughes (Uppsala University) for generously sharing a protocol for lambda red recombineering. We would like to also thank Duyen Bui and Kaili Zeng for their technical assistance.

